# Development of Cell-Derived Plasma Membrane Vesicles as a Nanoparticle Encapsulation and Delivery System

**DOI:** 10.1101/2023.08.06.552132

**Authors:** Mahsa Kheradmandi, Amir M. Farnoud, Monica M. Burdick

**Author notes:** Alpha Phase, Athens, OH 45701. To whom correspondence should be addressed: Monica M. Burdick, Ph.D.

## Abstract

**Background:** Developing non-invasive delivery platforms with a high level of structural and/or functional similarity to biological membranes is highly desirable to reduce toxicity and improve targeting capacity of nanoparticles. Numerous studies have investigated the impacts of physicochemical properties of engineered biomimetic nanoparticles on their interaction with cells, yet technical difficulties have led to the search for better biomimetics, including vesicles isolated directly from live cells. Cell-derived giant plasma membrane vesicles (GPMVs), in particular, offer a close approximation of the intact cell plasma membrane by maintaining the latter’s compositional complexity, protein positioning in a fluid-mosaic pattern, and physical and mechanical properties. Thus, to overcome technical barriers of prior nanoparticle delivery approaches, we aimed to develop a novel method using GPMVs to encapsulate a variety of engineered nanoparticles, then use these core-shell, nanoparticle-GPMV vesicle structures to deliver cargo to other cells.

**Results:** The GPMV system in this study was generated by chemically inducing vesiculation in A549 cells, a model human alveolar epithelial line. These cell-derived GPMVs retained encapsulated silica nanoparticles (50 nm diameter) for at least 48 hours at 37 °C. GPMVs showed nearly identical lipid and protein membrane profiles as the parental cell plasma membrane, with or without encapsulation of nanoparticles. Notably, GPMVs were readily endocytosed in the parental A549 cell line as well as the human monocytic THP-1 cell line. Higher cellular uptake levels were observed for GPMV-encapsulated nanoparticles compared to control groups, including free nanoparticles. Further, GPMVs delivered a variety of nanoparticles to parental cells with reduced cytotoxicity compared to free nanoparticles at concentrations that were otherwise significantly toxic.

**Conclusions:** We have introduced a novel technique to load nanoparticles within the cell plasma membrane during the GPMV vesiculation process. These GPMVs are capable of (a) encapsulating different types of nanoparticles (including larger and not highly-positively charged bodies that have been technically challenging cargoes) using a parental cell uptake technique, and (b) improving delivery of nanoparticles to cells without significant cytotoxicity. Ultimately, endogenous surface membrane proteins and lipids can optimize the physicochemical properties of cell membrane-derived vesicles, which could lead to highly effective cell membrane-based nanoparticle/drug delivery systems.

## Background

Liposomes are ideal for a wide variety of experimental and biological applications due to their unique physical properties that resemble biological membranes (1). For instance, antibody-functionalized liposomes have been used in our previous study to successfully localize to target cells (1). Nonetheless, in comparison to the complexity of the cell membrane, the arrangement of lipids and proteins is comparatively simplistic in manufactured systems, and the information gained from this model system is constrained by this limitation (2). Additionally, liposomes in their most basic forms are composed of only lipids and lack membrane proteins. Membrane proteins are critical features used by cells to control membrane thickness, transport nutrients, and control the exogenous interactions (3). While membrane protein incorporation into liposomes seems like a straightforward solution, it is not possible for all practical or desired purposes.

Further complicating matters, there is a severe lack of knowledge about the exact composition, position, order, and compartmentalization of proteins in the cellular plasma membrane (4). The plasma membrane is also dynamic - constantly rearranging, shifting, moving, and changing bilayer composition in response to environmental cues and intracellular circumstances. All told, it is difficult to mimic this narrow (5 to 10 nm), unstable, and complicated cellular compartment (5). Therefore, plasma membrane synthesis from scratch is not a practical technique to obtain an ideal biomimetic delivery model.

Cell membranes are an advanced source of materials for molecular delivery systems. Cell membrane-derived vesicles, much more so than synthetic lipid or polymeric nanoparticles, have a multicomponent characteristic that includes lipids, proteins, and carbohydrates (6). Due to their high functionalities and signaling platform similarities to the intact cell membrane, they can address various *in vivo* challenges, including the ability to minimize the immune system reaction (7). Moreover, the hollow-core structure of cell membrane-derived vesicles makes them a suitable coating material for different hydrophilic and hydrophobic pharmaceutical nanostructures. Cell membrane-derived vesicles have been used to coat a variety of therapeutic nanoparticles made of various materials and sizes, in addition to embedding small molecules within their exteriors (8).

Drugs encapsulated in cell membranes have also been shown to have a long-term, sustained release pattern. For the past three decades, red blood cells have been researched as delivery mechanisms for the ability to load and release various therapeutics, such as nucleic acids or synthetic drugs (9,10). Mishra et al. have monitored the release of doxorubicin from red blood cell vesicles; the release study showed more than half of the encapsulated drug was released slowly in the first 16 hours of incubation (11). Although several small molecules, such as doxorubicin (11), azidothymidine, and ethambutol (12) have been encapsulated in the core of erythrocytes, larger drug macromolecules and nanoparticles have proven challenging to encapsulate due to lower uptake and higher membrane disruption.

Cell membrane isolation and reconstruction is also another technique that has gained much attention during the last decade (13,14). However, isolation of plasma membrane from other cellular organelles is a complex and time-consuming process which requires the use of high pressures, sonication, strong chemicals, and/or detergents (15). Unsurprisingly, the disruption of functional membrane components, especially the folding of membrane proteins, has been reported during very long detergent or sonication-based membrane isolation techniques (16). Therefore, a new isolation technique would be greatly beneficial to obtain valuable information regarding the nanoparticle-plasma cell membrane interaction, without complicating interference from the endocytic process.

Giant plasma membrane vesicles (GPMVs) are micron-sized vesicles with the closest approximation to the size (<6 μm) and composition of the native plasma membrane. GPMVs hold promise for various applications, including research on coexisting fluid phases and membrane properties, which were previously performed on liposomes (2). Due to the high similarity with biological membranes, they are also promising candidates for drug encapsulation (including therapeutic nanoparticles) and delivery by (a) mitigating potential toxicity and (b) containing all membrane proteins, which could facilitate cellular uptake, targeting, and intracellular trafficking of a wide variety of cargos.

Trams et al. first identified cell membrane-derived, extracellular vesicles as delivery and intercellular communication means in 1981 (17). More recent investigations have shown that selected small molecules and amphiphilic quantum dots can translocate across the membrane into the lumen of GPMVs without the assistance of endocytic processes (18,19). In one of these studies, Säälik et al. demonstrated that GPMVs could encapsulate small cell membrane penetrable peptides by direct translocation across the plasma membrane (20). Another study by Pae et al. confirmed this finding while also suggesting that membrane proteins contribute significantly to the uptake of these small cell-penetrable peptides (18).

Phase segregation was reported after the direct membrane penetration method, leading the authors to conclude that certain cell-penetrable peptides could rearrange and change the lipid ordering of membranes (20). However, this same investigation indicated that uncharged, negatively charged, or weak-positively charged peptides and dextran could not penetrate into the vesicle lumen (20). Furthermore, (a) larger peptides with molecular mass of 2–3 kDa or higher and (b) low drug concentration solutions did not show any considerable uptake using the direct membrane penetration process (18,20). In another study, Silva et al. used serum-free media treatment for 48 hours to induce iron oxide loaded exosome production from 9 to 18 nm magnetic nanoparticles (21). However, it has been demonstrated that this slow vesiculation technique can lead to the altered protein composition within the final vesicle structure (22). Considering all of this, a new loading mechanism is required to provide a faster, efficient encapsulation mechanism for larger or not highly-positively charged entities.

The study herein introduces a novel nanoparticle loading mechanism using parental cell uptake before vesiculation. GPMVs were generated by chemically-induced vesiculation in A549 cells, a model alveolar epithelial cell line. After feeding fluorescent, negatively charged nanoparticles to cells, GPMV-based, core-shell, nanoparticle-vesicle structures were generated. This new GPMV system was evaluated for size, charge, stability, lipid and protein composition, and uptake and toxicity in two human cell lines, the parental A549 cells and model monocytic THP-1 cells.

## Results

### Size and Charge Characterization of GPMVs

The chemical induction method was used to produce a cell-derived, core-shell giant plasma membrane vesicle (GPMV) system in A549 human alveolar epithelial cells (23). The resulting blebs were imaged using wide field phase contrast microscopy, as well as confocal microscopy when using rhodamine-DOPE dye (**Fig. 1A** and **1B**). The average diameter of 115 representative GPMVs from 10 confocal images was determined to be 5.7 ± 2.3 μm, measured ImageJ software. Using the dynamic light scattering method, the average size of harvested GPMVs was measured as 4.9 ± 1.4 μm. There was no statistically significant difference between the sizes measured by the two techniques. Since the surface charge has been shown to have a significant impact on the biodistribution of delivery systems (24), laser Doppler anemometry was used to measure the surface charge of GPMVs. The surface charge of GPMVs was measured to be -29.4 ± 1.2 mV in PBS at 7.4 pH.

**Figure 1.**
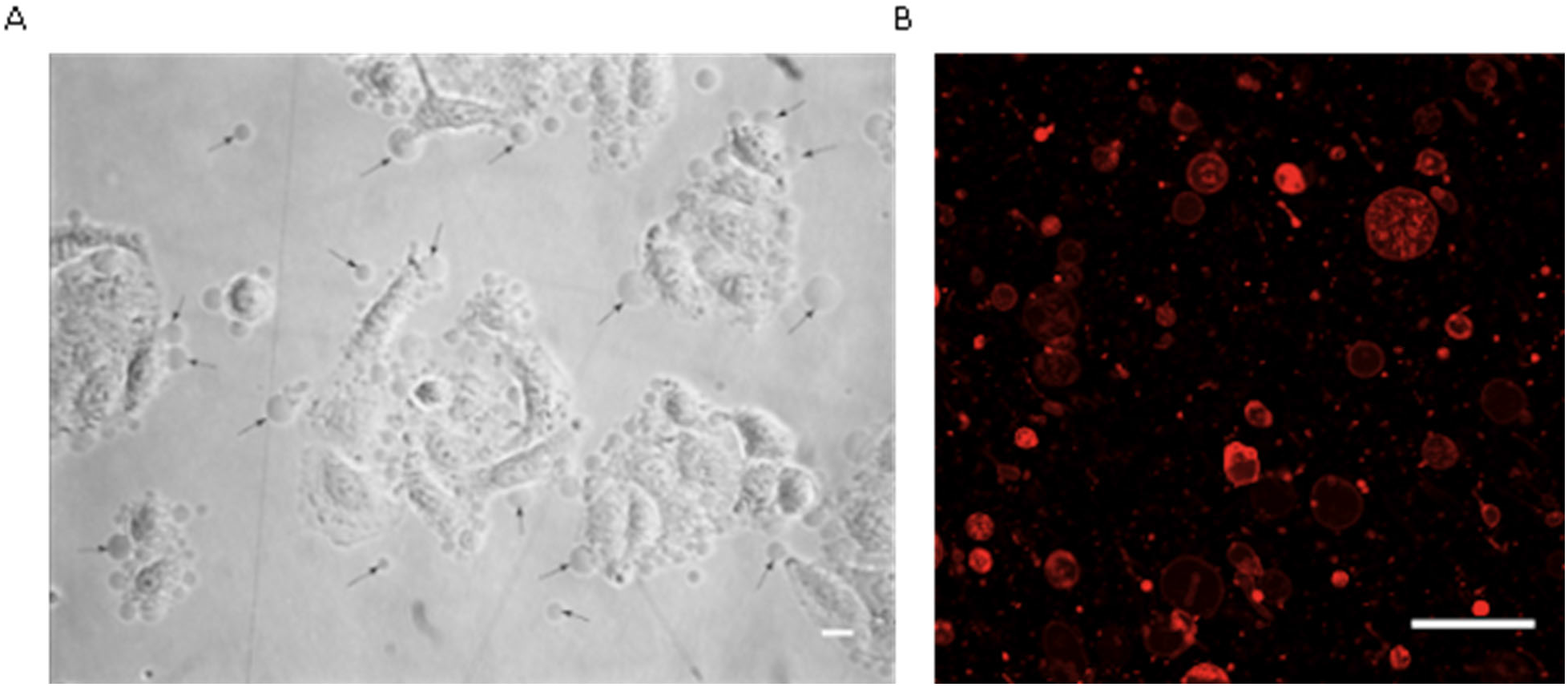
Visualization of GPMVs derived from A549 cells using (**A**) phase contrast microscopy (arrows indicate GPMVs next to parental cells) and (**B**) confocal microscopy imaging for Rhodamine (Red)-DOPE stained GPMVs. Samples were imaged at 100X magnification, and the scale bar is 5 μm. The experiments were repeated six times.

### Encapsulation of Nanoparticles in GPMVs

Herein, a new vesicle-based loading mechanism was introduced by incubating nanoparticles with parental cells. FITC-conjugated carboxyl-modified silica nanoparticles were first added to the culture media of A549 cells (i.e., parental cells) to uptake nanoparticles by endocytosis and diffusion. Then, chemical induction was used to generate nanoparticle-loaded GPMVs. Confocal imaging confirmed the presence of FITC-conjugated nanoparticles inside the Rhodamine-DOPE stained vesicles (**Fig. 2**). This implies that silica nanoparticles were successfully uptaken by A549 cells, followed by entrapment within vesicles that were excreted from the parental cells.

**Figure. 2.**
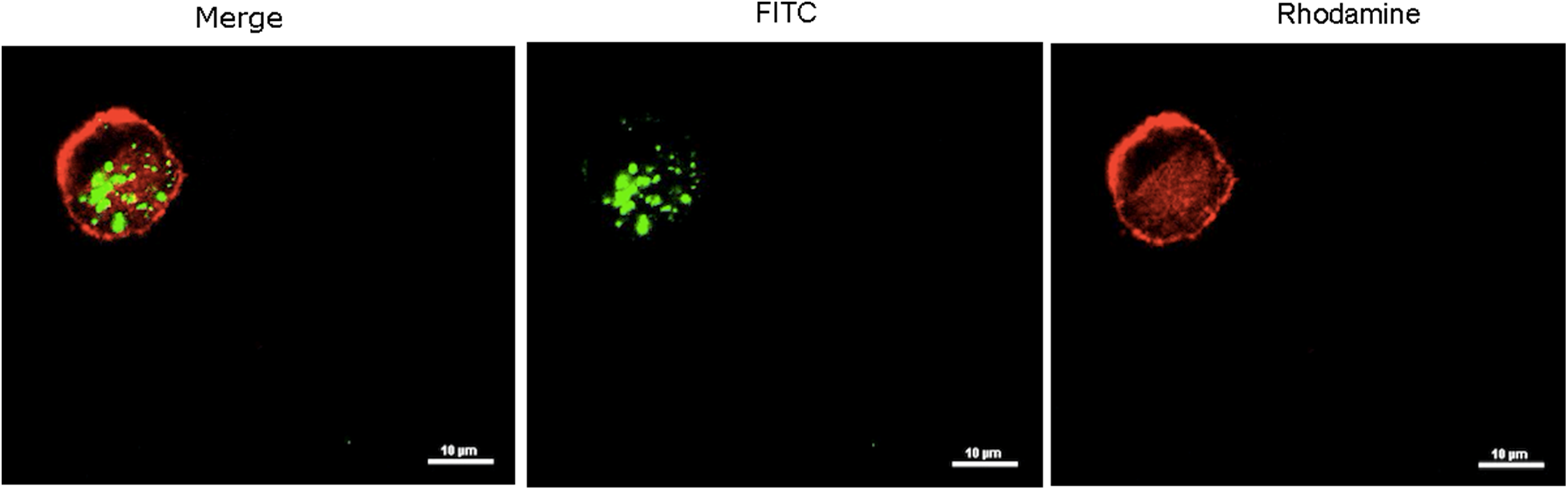
Internalized nanoparticles in A549 cell-derived GPMVs, examined by confocal fluorescence microscopy. GPMV membranes are red (Rhodamine-DOPE) and nanoparticles are green (FITC). Samples were imaged using a 100X objective, and the scale bar is 10 μm. The experiment was repeated six times.

### Stability Study of Nanoparticle-Encapsulated GPMVs

Stability testing is a major criterion to distinguish safe and reliable systems from undesirable fragile carriers. Thermal stability and integrity of FITC-conjugated carboxyl-modified silica nanoparticles in GPMVs from A549 cells were investigated for 48 hours in this study. Images clarified that nanoparticle encapsulating-GPMVs had spherical and intact shapes, showing that nanoparticles remained both encapsulated and stable inside the GPMV vesicles for at least 48 hours (**Fig. 3**).

**Figure 3.**
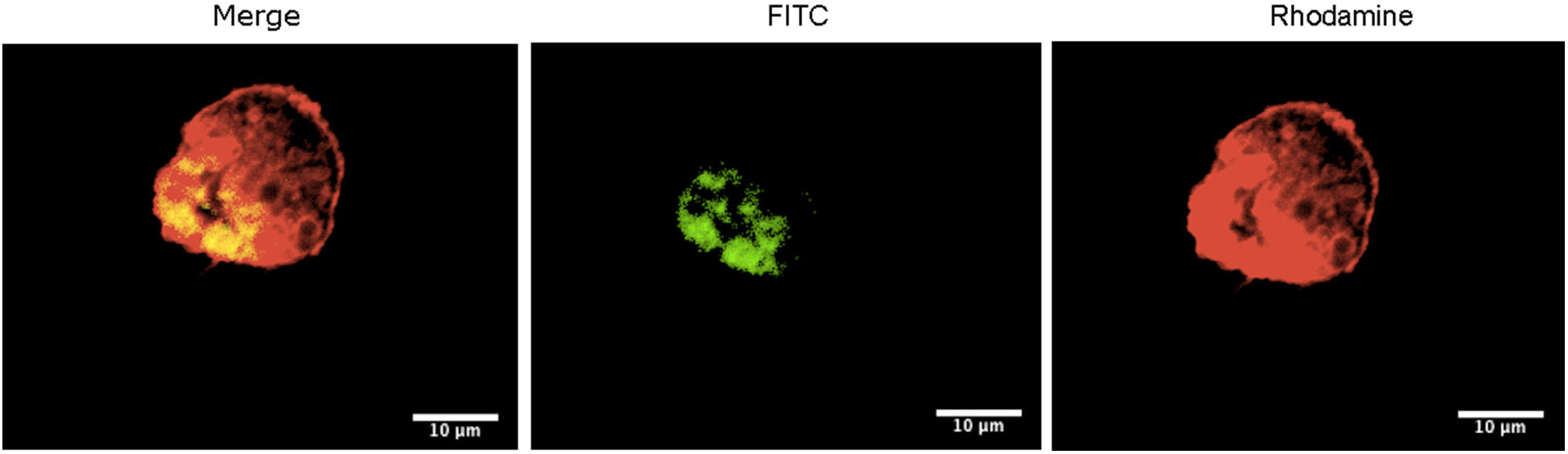
Stability study of nanoparticles encapsulated in GPMV after 48 hours as examined by confocal fluorescence microscopy. GPMV membrane and nanoparticles were visualized with rhodamine (red) and FITC (green), respectively. Samples were imaged at 100X magnification, and the scale bar is 10 μm. The experiment was repeated six times.

### Lipidomic and Proteomic Analysis of GPMVs

A comprehensive understanding of the nanoparticle-GPMV interaction is necessary for the clinical translation of *in vitro* assessments. Lipidomic and proteomic techniques provide important insights into biochemical responses of cells by identifying and quantifying numerous lipids and proteins (25). Lipid and protein profiling was applied here to investigate the potential impacts of encapsulating nanoparticles within the GPMV core-shell structure. A549 cells and carboxyl-modified silica nanoparticles were used to study the protein and lipid profiling of loaded- and unloaded-GPMVs. Vesicles showed the same lipid and protein profiles as reported for the cell plasma membrane with only minor changes in their lipidome and proteome, with or without encapsulation of nanoparticles. Lipidomics showed just one significant difference for ether-linked phosphatidylcholine (decrease in ePC level) in the lipidome of nanoparticle-containing GPMVs (**Fig. 4A**). Moreover, the protein abundance profile was consistent with the data reported by Eastlake et al. for normal retinae cells (**Fig. 4B**) (26). Nanoparticle-membrane interaction caused upregulated expressions in the glyceraldehyde-3-phosphate dehydrogenase (GAPDH) and human serum albumin (SPLAB) as compared to control unloaded GPMVs that may be caused by the intracellular nanoparticle corona. These results demonstrated that the lipid and protein composition of plasma membrane derived vesicles would not change drastically from exposure to silica nanomaterials.

**Figure 4.**
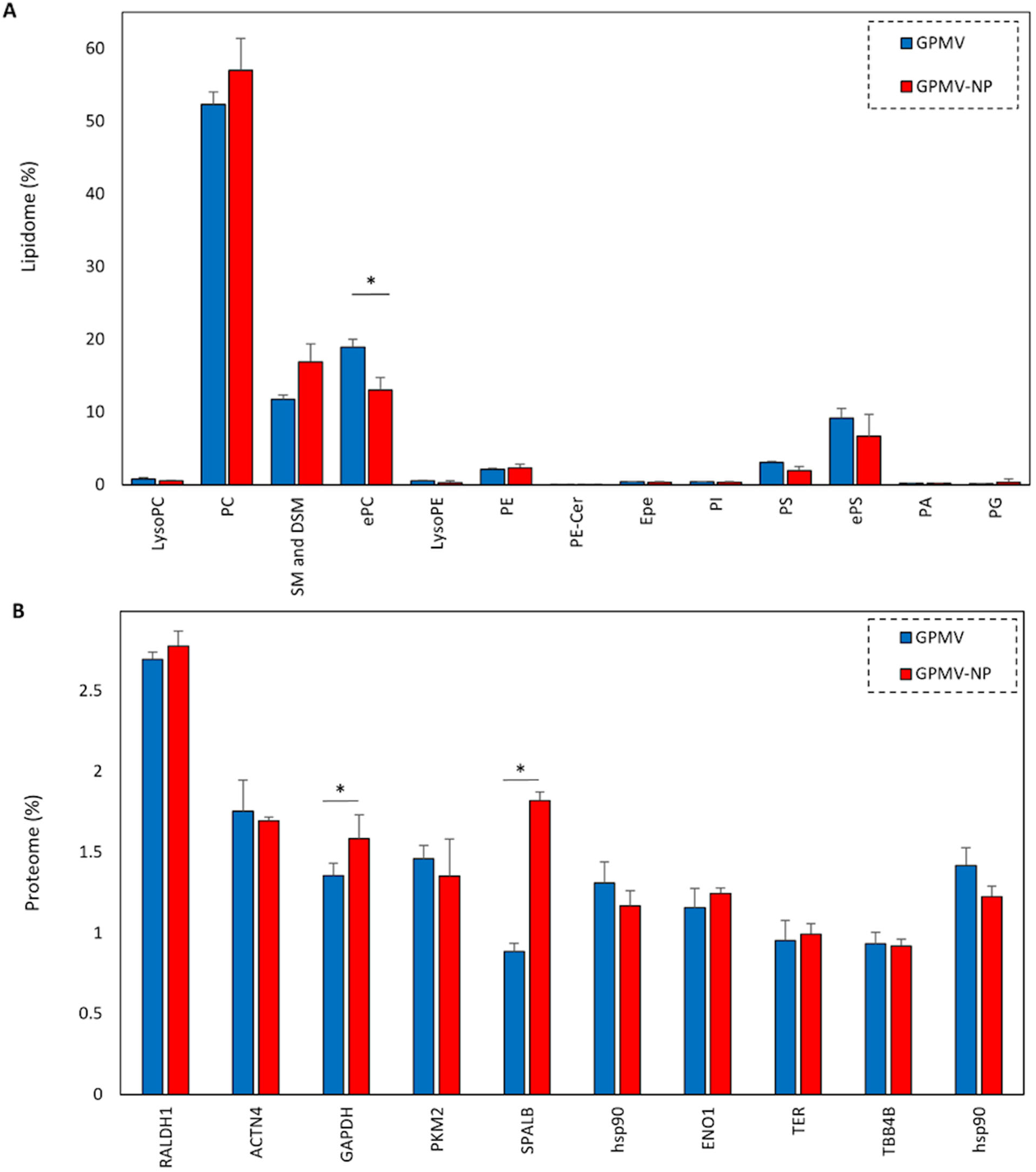
Analysis of lipid- and protein-profile changes in nanoparticles loaded and unloaded GPMVs. (A) Comparative lipidomics analysis between the 13 most abundant lipids. (B) Proteomic profiling of all membrane proteins. The experiment was repeated three times and error bars represent the standard deviation of three measurements. * indicates p < 0.01.

### Cellular Uptake and Release of Nanoparticle Loaded-GPMVs

Cellular uptake of large macromolecules and drug carriers is generally ineffective. As a result, developing efficient delivery systems for intracellular delivery of chemical and biological substances is desirable. Confocal imaging indicated that nanoparticle encapsulated in GPMVs were readily endocytosed in A549 (**Fig. 5A, top three rows**) and THP-1 cell lines (**Fig. 5A, bottom three rows**). As Fig. 5B indicates, a higher recipient cellular uptake level was observed for encapsulated nanoparticles (25.30 ± 5.43 for THP-1 and 25.21 ± 11.44 for A549 cells) compared to the other control groups, including the nanoparticles alone, GPMVs (not shown here), and control cells. Additionally, a higher level of control nanoparticles was uptaken with A459 cells (29.87 ± 3.94) compared to THP-1 cells (13.21 ± 10.74). This observation is consistent with previous findings that clathrin-mediated endocytosis is the main cellular uptake mechanism for 50 to 200 nm silica nanoparticles (27–29) and that A549 cells exhibit higher clathrin-mediated endocytosis compared to THP-1 cells (27). Therefore, it is implied that GPMVs can effectively facilitate delivering cargo to certain cells and can potentially serve as a novel nanoparticle/drug delivery system.

**Figure 5.**
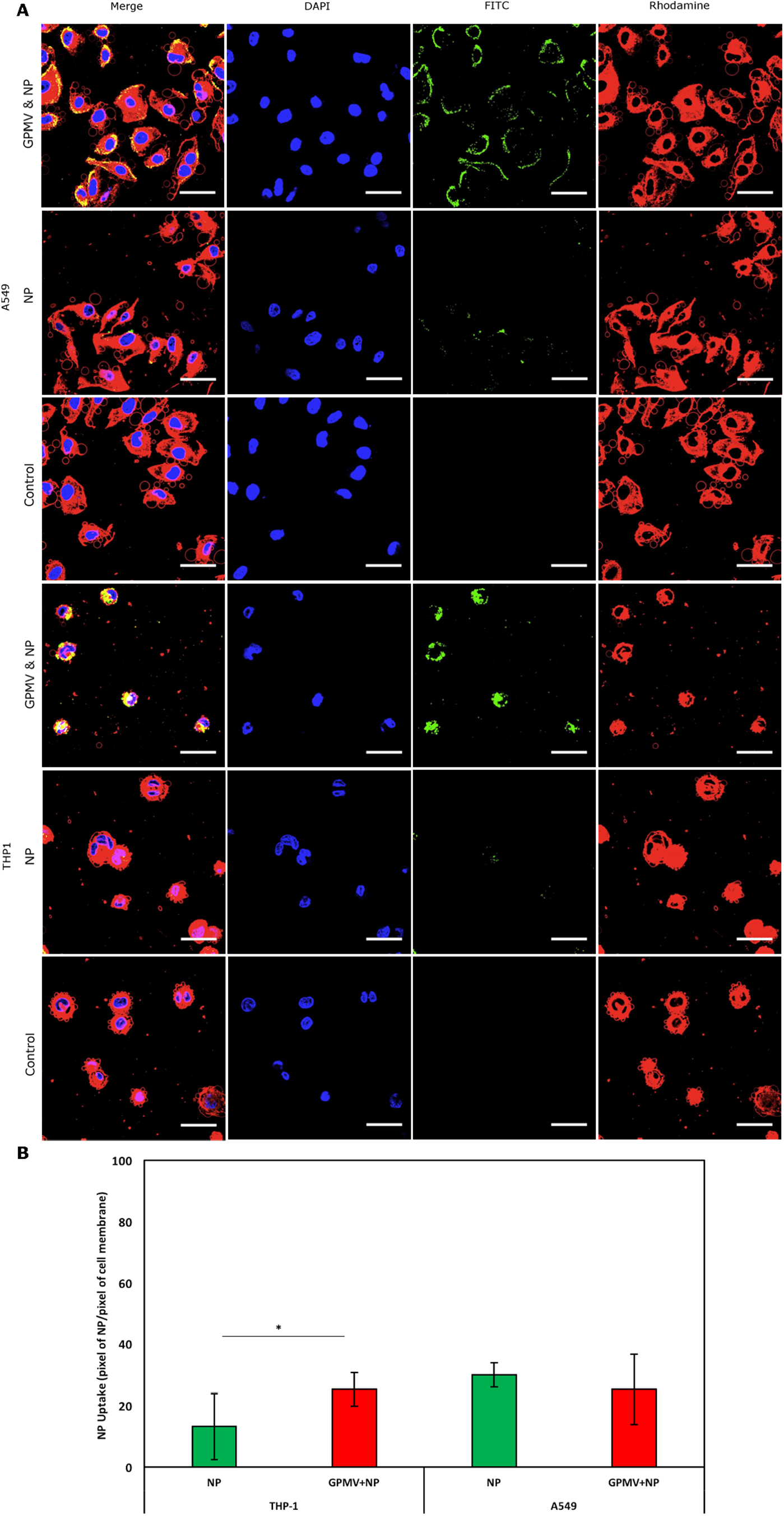
GPMV internalization in A549 and THP-1 cells. (A) Confocal imaging shows nanoparticle uptake before and after encapsulation within the GPMV vesicles. (B) Fluorescence intensity measurements using ImageJ software to quantify the number of green and red pixels, indicating relative expression of fluorescent nanoparticles and cell membrane areas. The plasma membrane and nucleus of adherent A549 cells were stained with rhodamine (red) and DAPI (blue), respectively, while silica nanoparticles (NP) were conjugated with FITC (Green). Four different groups were imaged for each cell type, including the nanoparticles, GPMV (not shown here), nanoparticle encapsulated GPMV, and control cells. Samples were imaged using a 60X objective, and the scale bars are 25 μm. The experiment was repeated three times. * indicates p < 0.01.

### Cytotoxicity Study of Nanoparticle Loaded and Unloaded GPMV Vesicles

The size and charge of plain, amine-, and carboxyl-modified silica nanoparticles were measured using dynamic light scattering and laser Doppler anemometry at 0.05 g/l in PBS at 7.4 pH (**Table 1**). The sizes measured using the dynamic light scattering technique were comparable with the reported nominal sizes of the nanoparticles (50 nm each).

**Table 1.** The hydrodynamic diameter and zeta potential of different surface-modified nanoparticles in PBS at 7.4 pH. The experiment was repeated three times.

| Sample | Size (nm) | Charge (mV) |
| --- | --- | --- |
| Plain Nanoparticle | $49.6 \pm 10.1$ | $-19.4 \pm 3.8$ |
| Amine Coated Nanoparticle | $50.1 \pm 10.5$ | $-18.5 \pm 1.4$ |
| Carboxyl Coated Nanoparticle | $52.1 \pm 8.4$ | $-16.7 \pm 1.6$ |

The cytotoxicity of silica nanoparticles encapsulated in cell membrane-derived carriers was first compared to control cell membrane vesicles alone and to free nanoparticles. Measured via the MTS assay, the control GPMV vesicles did not cause toxicity to cells, similar to the untreated control cells **(Fig. 6)**. Free carboxyl-modified silica nanoparticles did not show any significant toxicity either. Plain nanoparticles showed significant toxicity starting from 6 hours (p < 0.01). Amine-modified silica nanoparticles began to show significant toxicity after 24 hours (p < 0.1). Adding cell membrane-derived shells decreased the cellular toxicity in all groups, especially for plain nanoparticles (**Fig. 6**). These findings confirm that GPMV vesicles are able to deliver toxic nanoparticles with reduced cytotoxicity compared to free nanoparticles, even at concentrations that would otherwise be significantly toxic.

**Figure 6.**
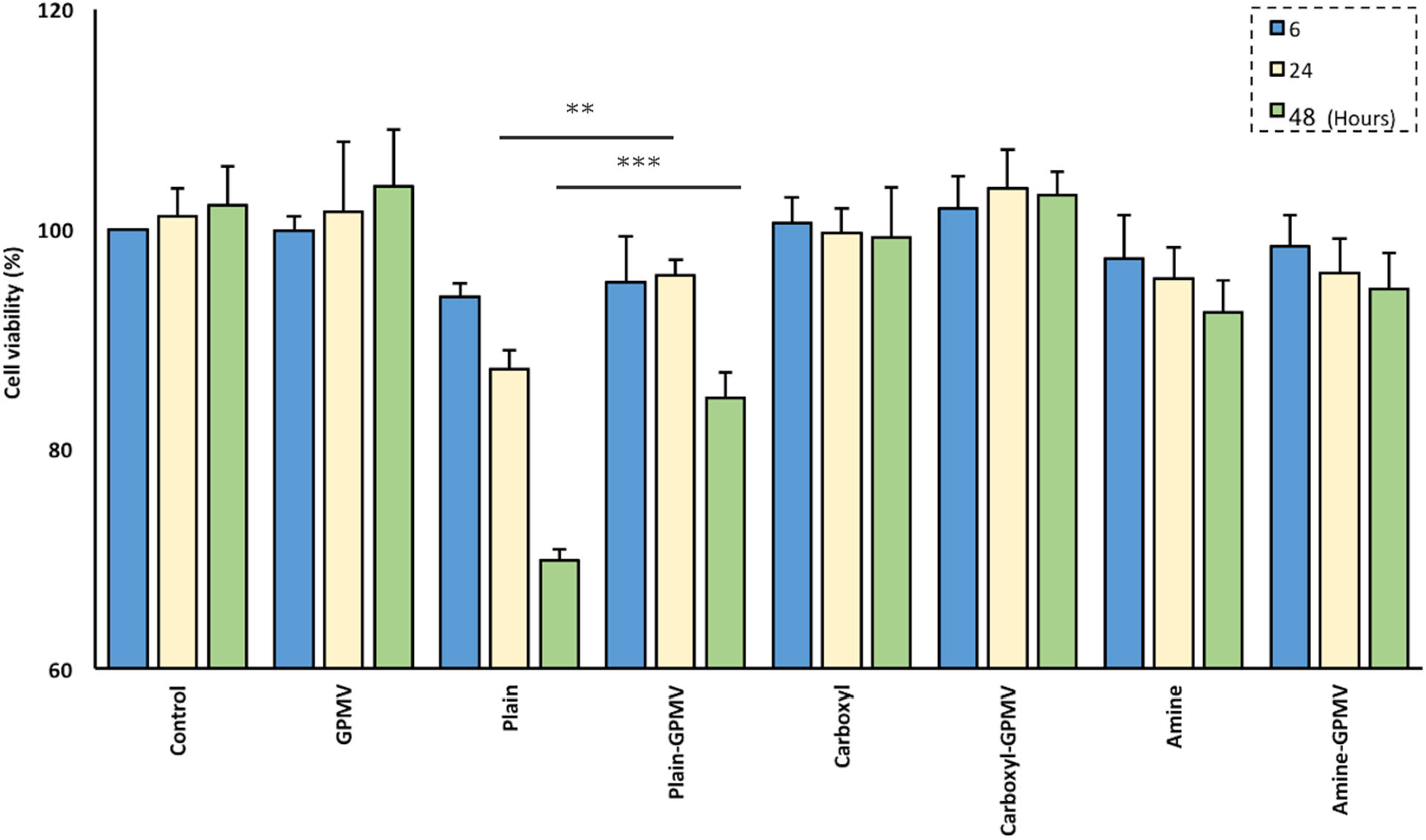
Effect of GPMVs and different nanoparticle-loaded GPMVs on A549 cell viability as examined by MTS assay at 6, 24, and 48 hours. Cytotoxicity induced by plain, carboxyl, and amine modified nanoparticles was measured by the MTS assay, which assesses cellular metabolic activities. The experiment was repeated three times, and error bars represent the standard deviation of the measurements. *** indicates p < 0.001 and ** indicates p < 0.01.

## Discussion

Nanomedicine is defined as the use of precisely engineered materials in the nanoscale for novel therapeutic and diagnostic modalities (30). Encapsulating nanomedicines within a biocompatible coating is a promising technique for resolving cytotoxicity, which is a major concern restricting the usage of several therapeutics (31). Different natural, artificial, and polymer-based delivery systems have been developed to provide safe passage, slow-release, and targeted delivery. While liposomes are the most widely used and well-studied nanocarriers for drug delivery (32), cell membrane-based systems have been demonstrated to offer better biocompatibility with little to no cytotoxicity compared to liposomes (6).

An exciting finding in previous reports revealed the possibility of loading highly positively charged peptides and cell penetrable compounds of low endogenous loading efficiency within GPMVs (18,20). However, this technique still excluded many potential therapeutic applications, including negatively charged nanoparticles (20). Herein, a novel loading mechanism for large silica nanoparticles (50 nm) in GPMVs has been introduced. Carboxyl modified silica nanoparticles were used for the loading purpose with the zeta potential of -16.7 mV, which is very close to the previously reported charge of -20.7 mV by Rocca et al (33). In contrast to the previous claim with Säälik et al. (20), these negatively charged nanoparticles have been successfully loaded in the core of cell membrane-derived shells (**Table 1**).

Parental cells (i.e, A549 cells) were incubated with negatively charged carboxyl modified nanoparticles. Then, GPMVs were vesiculated while entrapping the parental intracellular cytosol with 69.8 ± 21.1 % nanoparticle loading efficiency (**Fig. 2**). The nanoparticle entrapment yield is similar to the previously reported values (34,35). In one of these studies, Belwal et al. observed 71% loading efficiency for the encapsulation of silica nanoparticles inside liposomes composed of phosphatidylcholine and polyethylene glycol (PEG-2000) (34). In another study, Iehi et al. reported 71.4 % of silica nanoparticles inside whisker-formed Santa Barbara Amorphous-15 (SBA-15) nanoparticles (36).

This model was further characterized for membrane stability after encapsulation of nanoparticles. The stability of drug delivery systems refers to the extent that a container (e.g., vesicle) can retain cargo within a specified space and throughout its period of storage and use (37). Since leakage of delivery systems is an essential factor for the overall therapeutic index, we assessed the retention of nanoparticles encapsulated in GPMVs over 48 hours at 37 ^◦^C. While some fluctuation of the membrane was observed, confocal images indicated nanoparticles remained trapped inside the cell membrane-derived vesicles (**Fig. 3**). This result confirms the previously published leakage study by Säälik et al. (20), which indicated that at least for one hour after cell membrane penetrable peptides were loaded inside the GPMVs, entrapped peptides did not release significantly into the media.

Lipidomics and proteomic techniques have provided several important biochemical insights into nanoparticle alterations of membrane-derived vesicles. They can be used as early detection methods before most diseases or abnormalities enter terminal stages (38). Herein, we investigated any alteration in the lipid and protein profiling of cell membrane-based core-shell structure in response to the addition of nanoparticles. Our study showed only a few alterations between hundreds of lipid and protein components (**Fig. 4**). Therefore, the addition of carboxyl-coated silica nanoparticles to A549 cells did not cause considerable biochemical alternations in the cell membrane-derived vesicles.

Since several therapeutic paths are localized in subcellular compartments of recipient cells, it is critical to understand how drug carrier systems interact with desired cells and how the cargo-cell interaction affects cellular uptake and release (39). Furthermore, drug carriers’ uptake and final destination using different direct pathways or via endocytotic entry is often the main parameter to determine the overall therapeutic efficacy, kinetics, and transferability to animal model studies. It has been previously reported that GPMVs have the ability to fuse with target cell membranes, either directly with the plasma membrane or with the endosomal membrane following endocytic uptake (40). Even though GPMVs are relatively large (< 6 mm diameter, on the order of small cells) and that may limit applications, such functional behavior is encouraging and may bridge the realm between synthetic nanoscale strategies and new cellular therapies like CAR-T cells. Our confocal images indicated a strong uptake and uniform release of encapsulated nanoparticles inside the recipient A549 and THP-1 cells (**Fig. 5**). This positive finding suggests the high biocompatibility of the proposed system in future *in vivo* applications.

*In vitro* cytotoxicity assays often measure the ratio of cellular death in direct response to nanoparticle addition, which is an important criterion of new delivery systems. Different nanomaterials exhibit different cytotoxicity within the same recipient cells based on their particular physical, compositional, cellular uptake, and membrane interaction (41). Here, we have utilized three different plain, amine-, and carboxyl-modified silica nanoparticles to investigate the feasibility of using cell membrane-derived vesicles in reducing the cellular toxicity caused by these nanoparticles. MTS toxicity assay showed a different level of cell death in response to different nanoparticles. Also, GPMV vesicles were shown to reduce the toxicity caused by nanoparticles significantly in plain silica nanoparticle-loaded GPMVs, thereby enhancing A549 cell viability (**Fig. 6**) even though the uptake of NPs using GPMVs was similar to unencapsulated NPs (**Fig. 5B**). Our observation regarding the plain silica toxicity supports the findings of several other published articles (42,43). Fritsch-Decker et al. have suggested that silica nanoparticles damage the integrity of plasma membrane and cellular metabolic activities (44). Our results confirm previous reports that surface modification using amine and carboxyl groups significantly improves the biocompatibility of silica nanoparticles (40,45), as well as show a significant reduction in cellular cytotoxicity after encapsulating nanoparticles in cell membrane-derived vesicles (**Fig. 6**). More physiochemical studies are required to determine the effect of different surface functionalizations on the nanoparticle-cell membrane interactions before and after the vesiculation process and are the subject of ongoing investigations in our labs.

## Conclusions

The results of the present study clearly demonstrate that various compounds, namely large nanoparticles, can be loaded into GPMV vesicles to be used to transport therapeutics into the recipient cells. Furthermore, GPMVs may be used as favorable delivery systems with little to no cytotoxicity and high loading capacity due to their large (macro) sizes. Additionally, while GPMV vesicles lack the cortical cytoskeleton, ability to perform active endocytosis, and other metabolism-dependent biological processes, they still maintain the biochemical complexity of the native plasma membrane, including almost identical composition of lipids, proteins, and other matrix components, which distinguishes GPMV membrane models from other artificially made lipid vesicles and even live-cell membrane studies.

## Methods

### Materials

Promega CellTiter 96™ Aqueous One Solution Cell Proliferation Assay (MTS), HEPES buffer, KCl, CaCl_2_, and NaCl were purchased from Fisher Scientific (Pittsburgh, PA). Rhodamine-DOPE was purchased from Avanti Polar Lipids (Alabaster, AL). Sephadex G-25 in PD-10 desalting columns were purchased from GE Healthcare (Buckinghamshire, UK). Fluorescein isothiocyanate (FITC)-conjugated plain, carboxyl-, and amine-modified silica nanoparticles (50 nm) were purchased from Micromod Partikeltechnologie GMBH (Rostock, Germany). Dithiothreitol, chloroform, methanol, phosphate-buffered saline (PBS), paraformaldehyde, and other solvents were purchased from Sigma (St. Louis, MO). Human A549 alveolar lung carcinoma cells (as a model of aveolar epithelial cells) and human THP-1 acute monocytic leukemia cells (as a model of monocytes) were purchased from the American Type Culture Collection (Manassas, VA). Antifade mounting media with DAPI was purchased from Vector Laboratories (Burlingame, CA).

### Preparation and Characterization of GPMVs

RPMI 1640 supplemented with 10 volume percent fetal bovine serum (FBS) was used for A549 cell culture. GPMVs were generated as described previously by Levental et al. (23). In brief, confluent monolayers of adherent A549 cells in a 100 mm tissue culture dish were washed three times with PBS buffer and stained with 0.1 mg/ml rhodamine-DOPE for 10 minutes at room temperature. Then, cells were washed three times using the GPMV buffer containing 10 mM HEPES, 150 mM NaCl, and 2 mM CaCl_2_. After washing, cells were incubated with 27.6 mM paraformaldehyde and 1.9 mM dithiothreitol vesiculation agents at 37 °C for one hour under gentle stirring. Then, GPMVs were separated from the rich cellular supernatant with centrifugation at 500 RCF for 5 minutes. Also, two PBS washes following the centrifugation at 20,000 RCF for one hour at 4 °C were performed to separated GPMVs as a pellet at the bottom of the centrifuge tube. Blebs were imaged to investigate their size using confocal microscopy.

### Nanoparticle Encapsulation in GPMVs

To evaluate the ability of GPMVs to encapsulate intracellular content, we have added nanoparticles to the parent cells prior to the generation of GPMVs. Briefly, cells were first cultured at a 100 mm culture dish to reach a confluency of approximately 70% and pre-incubated with 0.05 g/l FITC conjugated carboxyl-modified silica nanoparticles (50 nm) for six hours, resulting in nanoparticle internalization, washed three times with PBS, and then chemically induced for GPMV generation in 27.6 mM paraformaldehyde and 1.9 mM dithiothreitol at 37 °C for one hour. GPMVs were separated as described above, with two PBS washes and two centrifugations. Confocal fluorescent imaging was used to confirm the encapsulation of nanoparticles inside the GPMVs.

### GPMV Membrane Integrity Study

The membrane integrity of GPMVs was evaluated using the stability assay of entrapped fluorescent nanoparticles. The stability of GPMVs at 37 °C was examined by monitoring the release of the fluorescent nanoparticles from their lumen. GPMVs were separated by centrifugation of GPMV-rich supernatant at 500 RCF for 5 minutes, washed three times with PBS, and harvested at 20,000 RCF for one hour and resuspended in the mounting media. Droplets of the nanoparticle-loaded GPMV solution were placed on a glass slide, incubated at 37 °C, and imaged using confocal microscopy after 1, 3, 12, 24, and 48 hours of incubation time.

### Quantitative Analysis of the GPMV Lipidome and Proteome Profile

Alterations in the proteomic and lipidomic profiles of GPMVs were investigated after the encapsulation of carboxyl-modified silica nanoparticles. A549 cells were cultured in 100 mm culture dishes to 70% confluency and chemically induced for vesiculation. Nanoparticle-loaded and unloaded GPMVs were harvested at 500 RCF for 5 minutes, washed three times at 20,000 RCF for one hour, and dried under vacuum in preparation for proteomic analysis at The Ohio State University Proteomics Facility (Columbus, OH).

Lipidomic samples were also prepared with an extra lipid separation step using the Bligh and Dyer method (46). Briefly, nanoparticle-loaded and unloaded GPMVs were mixed with 2.5:2.5:1.25:1 molar ratio of CHCl_3_:MeOH:water:sample followed by vortexing and centrifugation at 1000 RCF for 10 minutes. The organic phase was separated and dried under nitrogen gas for the mass spectrometry-based lipidomics at Kansas State University (Manhattan, KS).

### Membrane-Based Carriers for Nanoparticle Delivery Systems

The uptake of GPMVs, loaded with carboxyl-modified silica nanoparticles, in A549 and THP-1-derived macrophage cells, was observed using confocal imaging. A549 and THP-1 cells were both cultured in RPMI 1640 media with 10 volume percent FBS. Then, A549 and THP-1 cells were stained using 0.1 mg/ml rhodamine-DOPE for 10 minutes at room temperature. After the generation of nanoparticle-loaded GPMVs, 30 volume percent of the resulting vesicles from one 100 mm culture dish of parent cells were harvested as described before, resuspended in the cell culture media, and incubated in one well of 12-well culture plates for 6 hours with A549 and THP-1 cells lines, washed three times with PBS, and placed on the glass slides with DAPI-containing mounting medium for confocal fluorescent imaging.

### Cytotoxicity Study of GPMV Vesicles

The effect of GPMVs, nanoparticles, and nanoparticle-loaded GPMVs on cell viability was examined using the MTS cytotoxicity assay. Different nanoparticles were used, including plain, amine-, and carboxyl-modified silica nanoparticles. To this aim, A549 cells were grown to 70% confluency in a culture dish and incubated with nanoparticles, nanoparticle loaded-, and unloaded-GPMVs for 6, 24, and 48 hours. The nanoparticle binding and uptake efficiency were calculated 69.78 ± 21.08, as described below based on the average of three different nanoparticles; this yield was calculated based on the weight of the added nanoparticles compared to the samples after drying in the nanoparticle loaded-GPMVs.

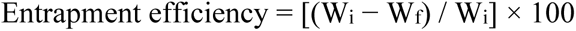

where W_i_ is the initial nanoparticle weight and W_f_ is the nanoparticle weight after loading in GPMVs. According to this nanoparticle loading yield, 69.78 volume percent of 0.05 g/l nanoparticle solution and 10 volume percent of loaded- and unloaded-GPMVs extracted from 100 mm culture dishes was added into each well of 96 well plates. Then, the MTS *in vitro* assay was used to measure cytotoxicity based on the colorimetric analysis of cellular metabolic activities at 490 nm. This cell-based toxicity experiment is a helpful tool to predict the potentially toxic effects of engineered membrane vesicles in cells.

### Statistical Analysis

All experiments in this study had at least three individual, independent replicates. One-way ANOVA with Dunnett’s test was performed to compare different samples. GraphPad Prism software was used for statistical analysis (La Jolla, CA). For each experiment, the average and standard deviation were reported in the mean ± standard deviation format. P-values of 0.05 were considered statistically significant.

## Declarations

### Ethics approval and consent to participate

Not applicable

### Consent for publication

Not applicable

### Availability of data and materials

All data generated or analyzed during this study are included in this article.

### Authors’ contributions

MK designed, performed, and analyzed experiments; interpreted data; and wrote and edited the manuscript. AMF designed and analyzed experiments, interpreted data, and edited the manuscript. MMB designed experiments, analyzed data, interpreted data, and wrote and edited the manuscript. All authors reviewed and approved the final manuscript.

## Acknowledgements

The authors gratefully acknowledge the use of the Ohio University Heritage College of Osteopathic Medicine Microscopy Core. We also thank J. Ayden Smith (Department of Chemical and Biomolecular Engineering, Ohio University) for assistance with manuscript preparation.

## References

1. Kheradmandi M, Ackers I, Burdick MM, Malgor R, Farnoud AM. Targeting dysfunctional vascular endothelial cells using immunoliposomes under flow conditions. Cellular and Molecular Bioengineering. 2020;13(3):189–99.

2. Jokhadar ŠZ, Klančnik U, Grundner M, Kebe TŠ, Hartman SV, Liović M, et al. GPMVs in variable physiological conditions: Could they be used for therapy delivery? BMC Biophysics. 2018;11(1):1.

3. Escribá PV, González-Ros JM, Goñi FM, Kinnunen PK, Vigh L, Sánchez-Magraner L, et al. Membranes: A meeting point for lipids, proteins and therapies. Journal of Cellular and Molecular Medicine. 2008;12(3):829–75.

4. Kersten GF, Crommelin DJ. Liposomes and ISCOMS. Vaccine. 2003;21(9–10):915–20.

5. Karp G. The structure and function of the plasma membrane. Cell and Molecular Biology: Concepts and Experiments, 3rd edn New York: John Wiley and Sons. 2002;122–82.

6. Tan S, Wu T, Zhang D, Zhang Z. Cell or cell membrane-based drug delivery systems. Theranostics. 2015;5(8):863.

7. Le QV, Lee J, Lee H, Shim G, Oh YK. Cell membrane-derived vesicles for delivery of therapeutic agents. Acta Pharmaceutica Sinica B. 2021;

8. Fang RH, Kroll AV, Gao W, Zhang L. Cell membrane coating nanotechnology. Advanced Materials. 2018;30(23):1706759.

9. Cinti C, Taranta M, Naldi I, Grimaldi S. Newly engineered magnetic erythrocytes for sustained and targeted delivery of anti-cancer therapeutic compounds. PloS One. 2011;6(2):e17132.

10. Harisa GI, Ibrahim MF, Alanazi F, Shazly GA. Engineering erythrocytes as a novel carrier for the targeted delivery of the anticancer drug paclitaxel. Saudi Pharmaceutical Journal. 2014;22(3):223–30.

11. Mishra PR, Jain NK. Folate conjugated doxorubicin-loaded membrane vesicles for improved cancer therapy. Drug Delivery. 2003;10(4):277–82.

12. Rossi L, Brandi G, Schiavano GF, Scarfi S, Millo E, Damonte G, et al. Heterodimer-loaded erythrocytes as bioreactors for slow delivery of the antiviral drug azidothymidine and the antimycobacterial drug ethambutol. AIDS research and human retroviruses. 1999;15(4):345–53.

13. Chen Z, Zhao P, Luo Z, Zheng M, Tian H, Gong P, et al. Cancer cell membrane–biomimetic nanoparticles for homologous-targeting dual-modal imaging and photothermal therapy. ACS Nano. 2016;10(11):10049–57.

14. Ran L, Lu B, Qiu H, Zhou G, Jiang J, Hu E, et al. Erythrocyte membrane-camouflaged nanoworms with on-demand antibiotic release for eradicating biofilms using near-infrared irradiation. Bioactive Materials. 2021;6(9):2956–68.

15. Lee YC, Bååth JA, Bastle RM, Bhattacharjee S, Cantoria MJ, Dornan M, et al. Impact of detergents on membrane protein complex isolation. Journal of Proteome Research. 2018;17(1):348–58.

16. Guo Y. Be cautious with crystal structures of membrane proteins or complexes prepared in detergents. Crystals. 2020;10(2):86.

17. Dang XT, Kavishka JM, Zhang DX, Pirisinu M, Le MT. Extracellular vesicles as an efficient and versatile system for drug delivery. Cells. 2020;9(10):2191.

18. Pae J, Säälik P, Liivamägi L, Lubenets D, Arukuusk P, Langel Ü, et al. Translocation of cell-penetrating peptides across the plasma membrane is controlled by cholesterol and microenvironment created by membranous proteins. Journal of Controlled Release. 2014;192:103–13.

19. Dubavik A, Sezgin E, Lesnyak V, Gaponik N, Schwille P, Eychmüller A. Penetration of amphiphilic quantum dots through model and cellular plasma membranes. ACS Nano. 2012;6(3):2150–6.

20. Säälik P, Niinep A, Pae J, Hansen M, Lubenets D, Langel Ü, et al. Penetration without cells: membrane translocation of cell-penetrating peptides in the model giant plasma membrane vesicles. Journal of Controlled Release. 2011;153(2):117–25.

21. Silva AKA, Di Corato R, Pellegrino T, Chat S, Pugliese G, Luciani N, et al. Cell-derived vesicles as a bioplatform for the encapsulation of theranostic nanomaterials. Nanoscale. 2013;5(23):11374–84.

22. Li J, Lee Y, Johansson HJ, Mäger I, Vader P, Nordin JZ, et al. Serum-free culture alters the quantity and protein composition of neuroblastoma-derived extracellular vesicles. Journal of Extracellular Vesicles. 2015;4(1):26883.

23. Levental KR, Levental I. Giant plasma membrane vesicles: models for understanding membrane organization. In: Current Topics in Membranes. Elsevier; 2015. p. 25–57.

24. Levchenko TS, Rammohan R, Lukyanov AN, Whiteman KR, Torchilin VP. Liposome clearance in mice: the effect of a separate and combined presence of surface charge and polymer coating. International Journal of Pharmaceutics. 2002;240(1–2):95–102.

25. Xu J, Bankov G, Kim M, Wretlind A, Lord J, Green R, et al. Integrated lipidomics and proteomics network analysis highlights lipid and immunity pathways associated with Alzheimer’s disease. Translational Neurodegeneration. 2020;9(1):1–15.

26. Eastlake K, Heywood WE, Banerjee P, Bliss E, Mills K, Khaw PT, et al. Comparative proteomic analysis of normal and gliotic PVR retina and contribution of Müller glia to this profile. Experimental Eye Research. 2018;177:197–207.

27. Hsiao IL, Gramatke AM, Joksimovic R, Sokolowski M, Gradzielski M, Haase A. Size and cell type dependent uptake of silica nanoparticles. Journal of Nanomedicine & Nanotechnology. 2014;5(6):1.

28. Blechinger J, Bauer AT, Torrano AA, Gorzelanny C, Bräuchle C, Schneider SW. Uptake kinetics and nanotoxicity of silica nanoparticles are cell type dependent. Small. 2013;9(23):3970–80.

29. Hu L, Mao Z, Zhang Y, Gao C. Influences of size of silica particles on the cellular endocytosis, exocytosis and cell activity of HepG2 cells. Journal of Nanoscience Letters. 2011;1.

30. Zhang L, Gu FX, Chan JM, Wang AZ, Langer RS, Farokhzad OC. Nanoparticles in medicine: therapeutic applications and developments. Clinical Pharmacology & Therapeutics. 2008;83(5):761–9.

31. Hung HI, Klein OJ, Peterson SW, Rokosh SR, Osseiran S, Nowell NH, et al. PLGA nanoparticle encapsulation reduces toxicity while retaining the therapeutic efficacy of EtNBS-PDT in vitro. Scientific Reports. 2016;6(1):1–13.

32. Sercombe L, Veerati T, Moheimani F, Wu SY, Sood AK, Hua S. Advances and challenges of liposome assisted drug delivery. Frontiers in Pharmacology. 2015;6:286.

33. Della Rocca J, Huxford RC, Comstock-Duggan E, Lin W. Polysilsesquioxane nanoparticles for targeted platin-based cancer chemotherapy by triggered release. Angewandte Chemie International Edition. 2011;50(44):10330–4.

34. Belwal VK, Singh KP. Nanosilica-supported liposome (protocells) as a drug vehicle for cancer therapy. International Journal of Nanomedicine. 2018;13(T-NANO 2014 Abstracts):125.

35. Montazeri M, Razzaghi-Abyaneh M, Nasrollahi SA, Maibach H, Nafisi S. Enhanced topical econazole antifungal efficacy by amine-functionalized silica nanoparticles. Bulletin of Materials Science. 2020;43(1):1–9.

36. Iehi AY, Shagholani H, Nikpay A, Ghorbani M, Soltani M. Synthesis and modification of crystalline SBA-15 nanowhiskers as a pH-sensitive metronidazole nanocarrier system. International Journal of Pharmaceutics. 2019;555:28–35.

37. Ahuja S, Dong M. Handbook of pharmaceutical analysis by HPLC. Elsevier; 2005.

38. Zhou Y, Wang H, Guo F, Si N, Brantner A, Yang J, et al. Integrated proteomics and lipidomics investigation of the mechanism underlying the neuroprotective effect of N-benzylhexadecanamide. Molecules. 2018;23(11):2929.

39. Garnacho C. Intracellular drug delivery: Mechanisms for cell entry. Current Pharmaceutical Design. 2016;22(9):1210–26.

40. Mulcahy LA, Pink RC, Carter DRF. Routes and mechanisms of extracellular vesicle uptake. Journal of Extracellular Vesicles. 2014;3(1):24641.

41. Jedrzejczak-Silicka M, Mijowska E. General cytotoxicity and its application in nanomaterial analysis. IntechOpen London, UK; 2018.

42. Chong CS, Cao M, Wong WW, Fischer KP, Addison WR, Kwon GS, et al. Enhancement of T helper type 1 immune responses against hepatitis B virus core antigen by PLGA nanoparticle vaccine delivery. Journal of Controlled Release. 2005;102(1):85–99.

43. Vranic S, Watanabe E, Miyakawa K, Takeuchi S, Osada Y, Ichihara S, et al. Impact of surface modification on cellular uptake and cytotoxicity of silica nanoparticles. 27 October 2020, PREPRINT (Version 1) available at Research Square https://doi.org/10.21203/rs.3.rs-96205/v1.

44. Fritsch-Decker S, An Z, Yan J, Hansjosten I, Al-Rawi M, Peravali R, et al. Silica Nanoparticles provoke cell death independent of p53 and BAX in human colon cancer cells. Nanomaterials. 2019;9(8):1172.

45. Bigdelou P. Role of Membrane Asymmetry in Nanoparticle-Erythrocyte Interactions [PhD Thesis]. Ohio University; 2020.

46. Bligh EG, Dyer WJ. A rapid method of total lipid extraction and purification. Canadian Journal of Biochemistry and Physiology. 1959;37(8):911–7.

